# Embedded Extremum Seeking: A Bio-synthetic Optimiser

**DOI:** 10.1101/2024.11.29.625997

**Authors:** Benjamin Smart, Lucia Marucci, Ludovic Renson

**Author notes:** L.M. and L.R. are co-last and co-corresponding authors.

## Abstract

In cybergentics, internal control uses synthetic genetic circuits to bypass the sensing and actuation limits of external control. Currently, embedded controllers are limited to basic controllers, like PID. Here, we introduce the Embedded Extremum Seeking Controller, the first example of a bio-synthetic optimiser. Extremum Seeking combines an integral and oscillator circuit to adjust system parameters and achieve a desired optimal steady state. This system uses an integral circuit and oscillator that already exist in synthetic biology. The Extremum Seeker is demonstrated *in silico* on a Labour Division system, where the controller tunes a kinematic parameter, optimising the ratio of the culture subpopulations to maximise production. This controller paves the way for embedded controllers that precisely direct cellular behaviour without needing external actuation and sensing.

## I. INTRODUCTION

Within cybergentics, external control is frequently used for cellular optimisation [1], [2], tuning the cellular environment to optimise production or proliferation. Unfortunately, external control is limited by its ability to sense and actuate [3], [4], [5]. Therefore, even the best optimisation algorithms struggle to externally optimise a cell culture [6].

Developing embedded controllers using synthetic genetic circuits can provide a system that self-regulates, bypassing these limitations. Thus far, Synthetic Biology has created circuits, such as toggle switches [7], oscillators [8], ‘concentration band-pass filters’ [9] and simple controllers, such as positive/negative auto-regulators [10], [11], lead/ lag controllers [12], an integral controller [13], [14], [15] and even a PID controller [16], [17].

Here, we introduce the Embedded Extremum Seeking Controller, a biosynthetic optimiser. This system uses an integral circuit [14], and an oscillator [18] to optimise a biological system’s steady state internally. Extremum Seeking, known since 1922 [19], has been applied to many different applications, including externally in bioreactors but not within cybergenetics [6]. It optimises a control input to achieve an optimal steady-state output in the system. ES is inherently model-free, not requiring a computational model within the controller to reach the optimum, making ES an ideal candidate for an embedded biological optimiser. In this paper, we have developed an ODE model that implements extremum seeking, built up of intracellular reactions that can be represented using mass action kinetics. The ES controller is used in the Labour Division’s case study.

Research shows that splitting different pathways between sub-populations of bacteria can reduce cellular stress when adding bio-synthetic circuits [20]. The optimal ratio between these sub-populations for maximum product output is unknown. We demonstrate, via simulation, how the embedded extremum seeker can optimise this population ratio.

This controller represents the first instance of embedded optimal control, paving the way for future embedded controllers that can precisely direct cellular behaviour without needing external sensing and actuation. This advancement could significantly boost production in industrial settings and lead to breakthroughs in medical treatments.

## II. Method

### A. Extremum Seeking

Extremum Seeking (ES) is not a new concept, with literature dating back to 1922 [19], developed to transfer the maximum power from overhead cables to trams. Yet over the last hundred years, ES has been used in a wide range of fields, from autonomous vehicles [21], drag reduction [22], combustion engines [23], jet engine stability [24], particle accelerators [25] and bio-processes [26], [27], [28]. However, it has only been externally implemented to maximise the bioreactor’s yield rather than internally using synthetic biology. In a system where an input can adjust the steady-state output, ES will seek the optimal input that maximizes the steadystate output.

ES works by modulating the input with an oscillating wave, producing the input, **u** and perturbing the steady state output, **y** as shown in Figure 1. The controller then uses this perturbation to estimate whether the output gradient moves towards or away from a maximum. The mean input is updated accordingly, via the averaging of the integrator, and the controller pushes the system towards the extremum.

**Fig 1.**
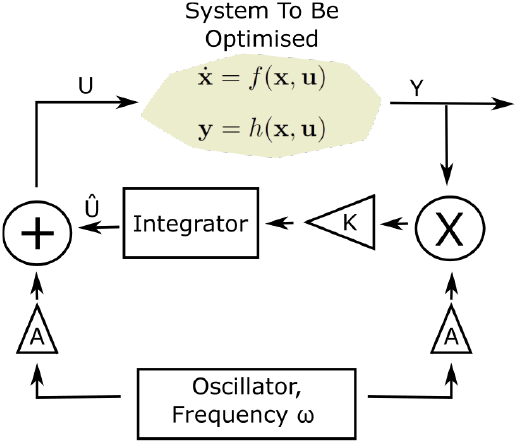
A block diagram displaying the three fundamental elements of an Extremum Seeker, the ‘system to be optimised’, the oscillator and the integrator. Including the three extremum seeking parameters, *K, A* and *ω*.

ES relies on separating three different time scales within the output signal. Importantly, no model of the ‘system to be optimised’ is required within the ES program. A High and Low pass filter helps reduce the time to reach the extremum but is non-essential [29]. It can be seen in Fig. 1 that three parameters are used to tune the ES. The amplitude gain *A*, the learning rate, *K*, and the oscillator frequency, *ω*.

The fastest dynamics in the response needs to be due to the ‘system to be optimised’, 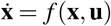, in Fig. 1 such that **y** = *h*(**x**) quickly reaches its steady state. Fig. 1 contains the model 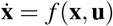 and **y** = *h*(**x, u**), however, this is merely to show that the controlled system can be controlled by **u** and the function is never known by the ES program.

The intermediate time scale is related to the frequency of the oscillator (or perturbation), *ω*, added to the input to explore a local neighbourhood of equilibrium around the current steady state, **y**_*p*_, caused by the addition of the oscillation *D* with a gain *A* such that **u** = **û** + *AD. ω* sets the time scale of the oscillator, controls the timescale separation of the ES and in turn controls the stability of the entire ES system, proved via Semi-global Practical Asymptotic(SPA) stability for this simple ES set up in [30]. Once *ω* is chosen, the gain of this oscillation, *A*, can be selected. As *A* increases, the convergence speed increases; however, the larger *A*, the ES will settle on a larger region surrounding the optimum. At a point, *A* becomes so large that there is a mismatch between the averaged system output estimate from the integrator and the actual system output. Therefore, increasing *A* should improve the convergence speed until a point in which the average from the integrator no longer matches the average of the actual system and the ES fails to reach the optimum [30].

Finally, the slowest time scale is the average change in the steady state of the output, also known as the learning dynamics [31]. At this time scale, the perturbations of the oscillator have been averaged out, and the controller implicitly estimates the gradient of the output, as shown in [19], achieved by the integrator within Figure 1. This updated input, **û**, is scaled by a gain *K*, the learning rate [19]. Similarly to *A*, as *K* is increased, the convergence speed will increase up until a point where the average from the integrator no longer matches the average of the actual system and the ES fails to reach the optimum [30].

Therefore, *ω* is initially chosen to achieve timescale separation from the ‘system to be optimised’, and then the amplitude of the oscillation *A* and the integral gain *K* are chosen to achieve the timescale separation between the integrator and oscillator, and can be fine-tuned for preferable dynamics, such as convergence speed. The ES will fail ω is too fast or *A* and *K* are too large not to achieve ti separation.

### B. Embedded Extremum Seeker Components

Biological systems typically contain time-varying parameters, usually modelled as large uncertainties or predicted parameter drifts. Model-based controllers need to adapt to these changes or incur a decrease in performance. However,

ES is model-free and has been shown to perform well on time-varying systems [32], [33], [34], managing to locate the extremum whilst retaining its stability guarantees [35], [36]. This makes ES a prime candidate for embedded control as it can deal with the time-varying nature of biological systems. ES consists of three fundamental blocks, as shown in Figure 1: A system to be optimised, an oscillator to cause perturbations and an integrator to average. Each of these blocks has to be represented by a biomolecular system.

#### 1) System To Be Optimised

The optimal concentration will be the ‘system to be optimised’ output. This system should reach a steady state faster than the oscillator and the integrator. We do not have to be able to record this steady state; it is just that the bacteria can sense the steady state. This system must contain one state or kinematic parameter that the integrator can act on and can be used to tune the output of interest and one output state that can act as the input to the integrator.

The embedded ES does not need a model of the ‘system to be optimised’; it only interfaces with this system’s living input and output. A model of each ‘system to be optimised’ used within this paper is only included for the simulations of the system response and will not be needed when the embedded ES is used *in vitro*.

#### 2) Integrator

A Bio-Synthetic integrator started to be developed in 2016 [37]. The integrator used here is taken from [14], with the system and response shown in Fig. 2A and the model

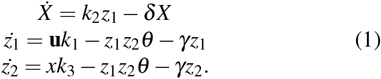

For the simulations, the integral parameters are [*k*_2_, *k*_3_, *θ, δ* , *γ*] = [1, 1, 0.01, 1, 0] producing the open loop response shown in Fig. 2B. It can be seen in Fig. 2 that the integrator is not perfect as *z*_2_ ≠ ∫ **u***dt*, yet it does carry some integral action, increasing when **u** is constant and remaining constant when **u** = 0.

**Fig 2.**
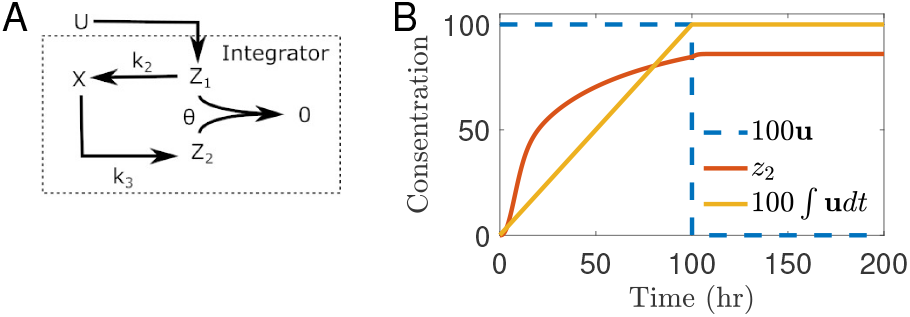
A) The molecular interactions that make up the integrator. B) The integrator’s open loop response to the step input is shown with the parameters discussed by Eq. (1) and *k*_1_ = 1.

#### 3) Oscillator

The Repressilator is used here [18], containing three cytoplasmic proteins that oscillate, *D*_1_ to *D*_3_,

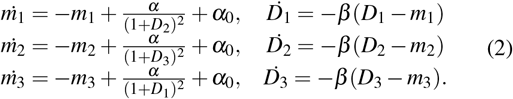

Each protein inhibits the production of the mRNA associated with the next protein along the chain, as shown in Fig. 3A. The nonlinearity in the system causes the oscillation due to the delay caused by transcription and translation. For the simulations, the Repressilator parameters are [*α, β, n, α*_0_] = [112, 3, 2, 10^−3^*α*] following the marked ‘X’ in [18], following the free response shown in Fig. 3B.

**Fig 3.**
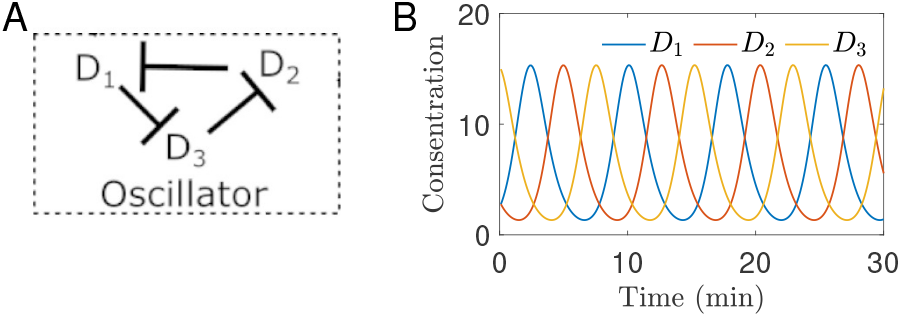
A) The molecular interactions that make up the Repressilator oscillator. B) The free response of the Repressilator.

### C. Integral and Oscillator Selection

The examples of the integrator and oscillator models used here are chosen with respect to the dynamics of the ‘system to be optimised’. It is stressed that extremum seeking will stably reach the optimum as long as there is timescale separation between the three fundamental blocks shown in Fig. 1, as described in Section II-A. ES may still reach the optimum without time scale separation, but there is no guarantee, and the theoretical stability guarantees cannot be applied [30]. Therefore, once the ‘system to be optimised’ has been identified, an oscillator with an appropriate frequency, *ω*, can be identified, and the integrator structure/parameters can be chosen.

#### 1) Integrator

The integrator has to be the slowest part of the ES system. As discussed in [15], the parameter *δ* can be increased, and *k*_2_ or *θ* within Eq. (1) can be decreased to slow the integrator’s response time. Therefore, *θ* has been reduced to the smallest value of the allowable range discussed in [15].

As long as the system implements integral action (or perfect adaptation), the exact structure of the system is flexible within the ES controller. We have achieved successful ES with the three state integrator in Eq. (1) and the four state structure discussed in [37]. In an attempt to reduce the number of states in the model and molecules required to be grown by the cell, the three state model is presented here.

#### 2) Oscillator

An oscillator with the appropriate timescale should be chosen [38] such that it seems to oscillate the steady state of the ‘system to be optimised’ rather than oscillating the transient response, and therefore the central timescale of the three step timescale separation. The oscillator needs to perturb the integral state both positively and negatively. This is initially difficult with only positive concentrations. To explore both an increase and decrease in the ES input, *U* is perturbed by the difference between two out-of-phase concentrations, *D*_1_ and *D*_2_, where *D*_1_ − *D*_2_ oscillates approximately around zero. Other oscillator setups that successfully add both positive and negative perturbations could also be used. The oscillator states perturb the states of the integrator, but the oscillator dynamics are decoupled from the rest of the ES setup. Therefore, the observed dynamics of the oscillator should not be significantly affected by adding it to the ES.

We have achieved successful ES with both the Hasty oscillator [39] and Repressilator [18], but the Repressilator was chosen for its faster frequency, *ω*, and near zero DC gain of *D*_1_ −*D*_2_ in Eq. (2).

### D. Embedded Extremum Seeker

Within biomolecular systems, it has been shown that degradation [40], transcription [41], translation [42], annihilation [13], enzyme and activation reactions can be modelled as proportional to the concentration of the reactant species, within a suitable range [42]. Within all the models discussed, only proportional promotion (displayed as →), proportional inhibition (shown as ⊣) and Hill functions 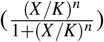 [43], are used as ODE terms to represent biomolecular reactions. From a schematic, a deterministic model is produced with one ODE for each species within a network and an ODE term for each reaction related to that species.

The biological blocks discussed in Section. II-B can be connected to form an ES. Fig. 4 shows the reactions needed to connect the oscillator to the integrator, implementing the addition and multiplication boxes in Fig. 1 to form an ES. Out-of-phase promotion/inhibition reactions, inspired by the dynamics of the Hasty oscillator [39], are needed to both increase and decrease the input to form the addition block within the integrated state, *z*_2_ and the multiplication block, along with the output of the ‘system to be optimised’ on *z*_1_. These addition and multiplication blocks in Fig. 1 can be seen by the four added ODE terms,

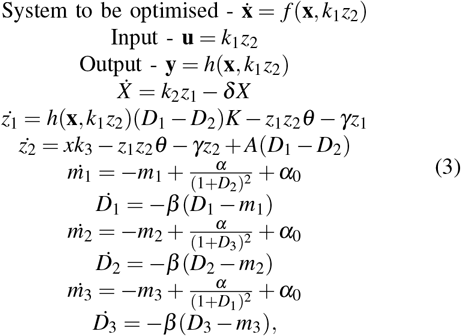

forming the embedded extremum seeker shown in Fig. 4.

**Fig 4.**
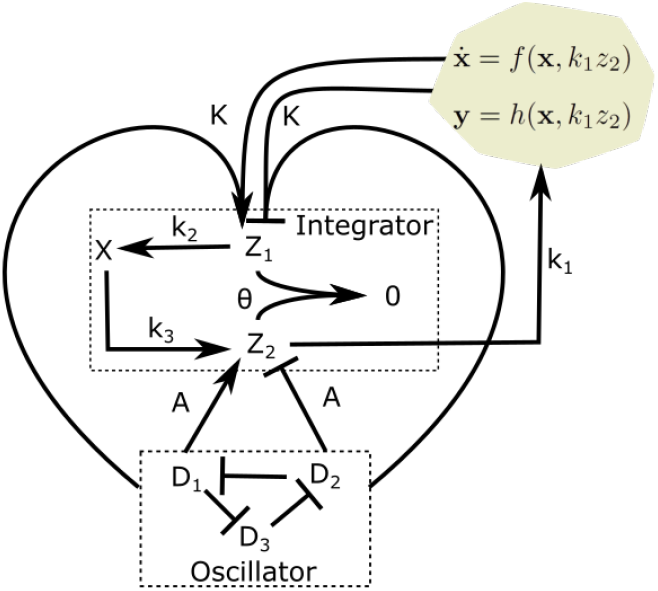
The biological network forming the ES, as written in Eq. (3). Including the three ES parameters, *k*_1_, *A* and *K*.

This ES model contains three added mass action kinetics: *A, K* and *k*_1_. *A* and *K* are the two ES parameters in Fig. 1, equivalent to the amplitude gain and learning rate, respectively, as discussed in Section II-A. However, *k*_1_ describes a phenomenon unique to the embedded ES. A computational integrator can calculate the integral of an input over a large range of magnitudes as they follow a mathematical formula. Whereas the integrator used here is a living bio-synthetic system, being produced in response to the input. *k*_1_ is trying to make up for the inflexibility in the working range of the bio-synthetic integrator. Due to production rate constraints, there is a feasible range in which the bio-synthetic integrator can operate. Therefore, when using the ES, *k*_1_ will need to be tuned as the user finds this suitable range of the integrator to effectively act on the ‘system to be optimised’.

### E. Biological Implementation

The Repressilator in [18], uses the three repressor proteins, lacI, tetR and cI. The protein, cI, is produced from *λ* phage, which is used in the Repressilator to repress; however, it has dual activity [44] so that its promoter site can be used for both promotion and repression and therefore would become *D*_1_ in Fig. 4, repressing tetR (*D*_3_) and promoting *z*_1_ and *z*_2_. LacI can repress *z*_1_ and *z*_2_ and repress cI, forming *D*_2_.

The integrator can be implemented with either transcriptional regulation [13], using sense/antisense transcription factors for *z*_1_ and *z*_2_ or translational regulation [45], [14] using mRNA and sRNA annihilation pairs for *z*_1_ and *z*_2_. The cI and lacI proteins will be transcription factors for *z*_1_ and *z*_2_ and, therefore, could directly affect the production rate of a translational level integrator. Otherwise, the cI and lacI would need to produce the sense/antisense transcription factors for *z*_1_ and *z*_2_, adding a time delay related to both the transcription and translation of *z*_1_ and *z*_2_, and the potentially significant amino acid resources and ribosome usage. Therefore, the integrator should be implemented as in [45] where *X* forms the ECF32 protein and *z*_1_ and *z*_2_ are corresponding mRNA and sRNA.

The biological realisation of the embedded ES system should be seen assimilar to the computational ES: modular. The oscillator and integrator blocks could be split and placed within separate cell strains if needed, as long as the output of the ‘system to be optimised’ affects *z*_1_ and can be tuned by changes in *z*_2_.

## III. Results

### A. Static Map

The stability and performance of the ES are characterised by the time scale separation of the three elements of the controller, as discussed in Section II-A. The controller has been used on a static map as a first attempt. This has no practical implementation as the static map can not be implemented within the bacteria. Still, it highlights the potential of this embedded ES controller when the timescale separation of the system to be optimised is guaranteed. Allowing performance and sensitivity analysis of just the integrator and oscillator combination.

#### 1) Simulation Results

The static map, 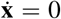 and *y* = *h*(**u**) = (25 (5 (**u**))^2^)*/*10 has been added to the system described in Eq. (3) within Fig. 5.

**Fig 5.**
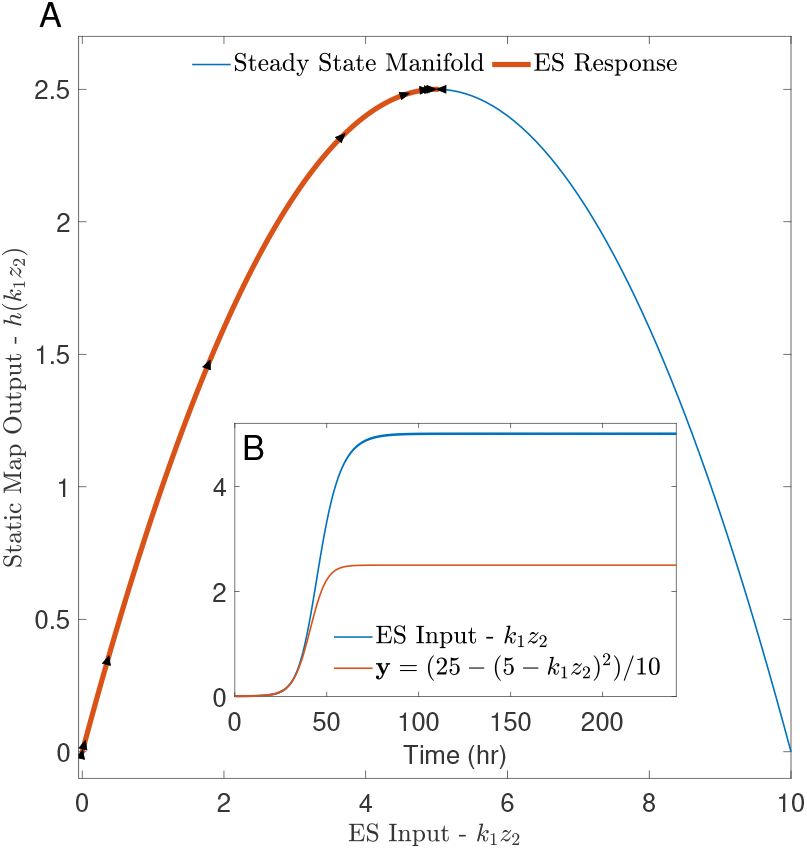
The ES controller response to a static map ‘system to be optimised’. A) The ES response, **y**, compared to the equilibrium manifold, **y** = *h*(**u**) = (25− (5− (**u**))^2^)*/*10 with an arrow every ten hours in the direction of the ES response. B) The input signal, **u** = *k*_1_*z*_2_, acting on the static map and the resulting output signal, **y** = *h*(*k*_1_*z*_2_) = (25− (5 − (*k*_1_*z*_2_))^2^)*/*10 in time. ES parameters: *k*_1_ = 0.25, *A* = 0.1 and *K* = 25.

It can be seen in Fig. 5 that the embedded ES manages to reach the optimum of the static map. Initially, the ES has a slow increase of the input, *u*, as the integrator passes through its transient response. Subsequently there is a large push from the ES controller to increase the input as it senses a large gradient. Once the ES reaches the maximum, it settles on an oscillation about the maximum, as can be seen by the change of arrow direction about the peak of the input/output plot. The filter takes 55 hours to reach the maximum. After this, the input and output continue to oscillate around the maximum as the perturbations from the oscillator are still added to the system.

This ‘system to be optimised’ reaches a steady state instantly, the oscillator has a time period of roughly 8 minutes, and the integrator has been tuned to act slowly according to [15]. Therefore, there is a successful timescale separation between the static map and the oscillator/integrator combination.

#### 2) Sensitivity Analysis

The parameters *k*_1_, *A* and *K* are chosen to tune the ES, but it needs to reach the maximum for a range of these parameter values. The speed and dynamics in achieving the maximum can vary between parameter sets within this range, but the controller should robustly reach the maximum.

Fig. 6 displays the steady state of the ES system (not the open loop of the ‘system to be optimised’ but the closed loop ES system in Fig. 4). It can be seen that within the parameter ranges 0.1 → *k*_1_ → 0.4, 0 *< A* ≤ 80 and 0 *< K* ≤350, the ES controller manages to reach the optimum of 2.5 *±* 10%.

**Fig 6.**
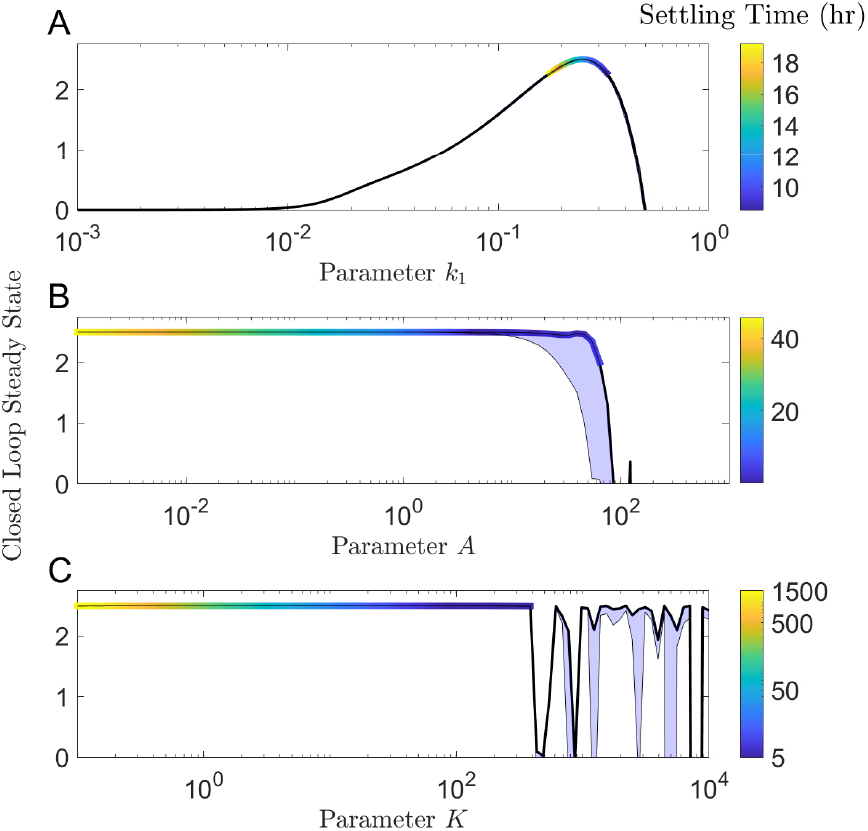
The Range of the ES parameters that achieve steady state. For each parameter, the mean steady state is surrounded by the amplitude of its steady state oscillation, which is displayed as the shaded blue area. For the range of parameters that successfully reach the optimum steady state, the colour of the mean steady-state line varies with the time taken to reach the steady state. A) The range of parameter *K*. B) The range of parameter A.C) The range of parameter *k*_1_. When not varied, *k*_1_ = 0.25, *A* = 0.1 and *K* = 25.

It can be seen in Fig. 6A that *k*_1_ does not have a wide feasible range, reaching 2.5 ±10% between only 0.1 →*k*_1_ → 0.4. Although the feasible operating range of the integrator limits the embedded ES, as discussed in section II-D, it can be seen that for over 1.5 orders of magnitude, *k*_1_ improves the output of the static map. For practical implementations, this would still have significant improvements to biological systems.

Parameter *A* controls the perturbation amplitude added to the input state. Fig. 6B displays how these input perturbations affect the system’s output. It can be seen that, as described in Section II-A, as *A* increases, the settling time decreases (the line gets darker) until a point in which the ES fails to reach the optimum at *A* = 100. The oscillations about the optimum are very small until *A* = 10, at which point the ES will visibly oscillate about the maximum; it can be seen that as *A* increases further, the ES exists in a larger neighbourhood of the optimum, also as expected.

Fig. 6C shows that *K* controls how aggressively the controller works towards the optimum, known as the learning rate. It can be seen that the smaller *K* is, the slower the settling time. However, as *K* increases, there comes the point where *K* becomes too large and the time scale separation between the oscillator and the integrator is no longer achieved, as described in Section II-A, and the controller response becomes unstable and unpredictable.

#### 3) Fano Factor

The Fano Factor is used to quantify the sensitivity of the steady state of the output,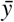 to the ES parameter changes. One thousand simulations are carried out where all parameters are held constant between simulations, apart from the parameter of interest, *P*, which is drawn from the normal distribution, *P*^*^ ∼ *𝒩* (*P*, 0.3*P*), each simulation. The Fano Factor,

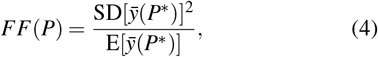

displaying the effect of a 30% uncertainty in parameter *P* on the steady state of the output.

It can be seen in Table. I that the Fano Factor of each ES parameter *A* and *k*_1_ are very small, showing that the controller provides robust optimisation for a wide range of

**TABLE I.** The Fano Factor, *FF* IN EQ. (4), FOR EACH PARAMETER, *P*.

| $P$ | $k_1$ | $A$ | $K$ | $k_1 z_2$ |
| --- | --- | --- | --- | --- |
| $FF(P)$ | 0.039 | $6.92 \times 10^{-10}$ | $4.18 \times 10^{-5}$ | 0.057 |

ES parameter values, as shown in Fig. 6. To compare the sensitivity of the ES to changes in its internal parameters to the sensitivity of the static map, **y** = *h*(*k*_1_*z*_2_) = (25 − (5 − (*k*_1_*z*_2_))^2^)*/*10, to changes in the input, **u** = *k*_1_*z*_2_, the Fano Factor of the input, *FF*(*k*_1_*z*_2_), is also included in Table. I. *FF*(*k*_1_*z*_2_) = 0.057, emphasising how insensitive the optimum is to changes, *A* and *K*, by over three orders of magnitude. However, it can be seen that the system is most susceptible to changes in *k*_1_, as seen in Fig. 6A, showing that for this static map example, changes in *k*_1_ restricted the performance of the ES controller as it struggles to make up for changes in *k*_1_ by changing the integral state *z*_2_, as discussed in Section IV.

### B. Labour Division

Here, we used ES to optimise the population ratio between two consortium cell strains. The optimal ratio between these sub-populations for maximum product output is unknown. The initial ratio of subpopulations can be chosen, but there is currently limited control of the populations online whilst only the final product concentration is observed, and the initial ratio of the next batch is slowly iterated over each run. The embedded ES could optimise this population ratio online within the first bioreactor batch.

Within synthetic biology, the genetic circuits that code for these functional systems are usually coded into one plasmid, grown in one bacteria population [46], [47]. It has been shown that the added metabolic burden increases cellular stress, forming an evolutionary pressure to drop the plasmid. Due to this competition, subpopulations form within the colony, competing with each other, increasing the rate at which the engineered plasmid is dropped from the population [20].

It has been shown that splitting different pathways between sub-populations of bacteria can both decrease cellular stress and increase total production when adding bio-synthetic circuits [20]. These microbial communities interact with cross-feeding mutualism [48], where each sub-population needs a product to survive that they cannot produce but obtain from another sub-population. This division of labour makes the production system more robust and constant as there is less competition between subpopulations. Therefore, many engineered cell systems are turning from one-strain consortiums to multi-cell consortiums. This ES can maximise production by optimising the cell strain population ratios.

#### 1) Model

The labour division system requires one mass action parameter or cellular concentration that can be used to tune the population size of both strains of the two-strain consortium and a concentration to maximise by optimising this population ratio.

Fig. 7 displays the two-strain model containing two bacteriocin molecules, *B*_1_ and *B*_2_, used to stabilise each cell strain subpopulation, *N*_1_ and *N*_2_, respectively, living of a shared metabolite store, *s*. One quorum sensing molecule is used to inhibit the action of both bacteriocin, as developed in [49]. The quorum sensing molecule simultaneously affects both cell populations and can be used to tune the cell strain population ratio. This model does not contain the production of a product. Therefore, we adapt the model from [49] to include the production of Resveratrol, with *N*_1_ producing a metabolite needed by *N*_2_ to produce the Resveratrol, *R* [20]. The system in Fig. 7 forms the model,

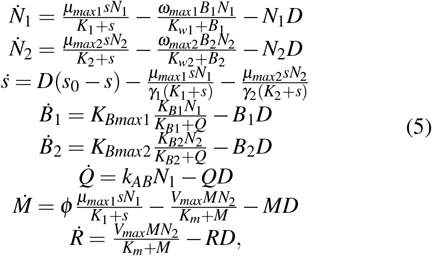

using parameters within the ranges discussed in [49], [20],

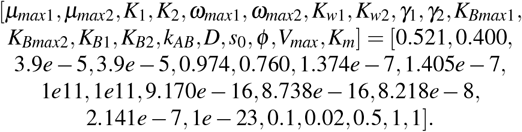

This model forms the ‘system to be optimised’ within the ES, using *k*_*AB*_ coupled to the input as 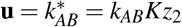, to control the concentration of the quorum sensing molecule, *Q*, tuning the population ration. When 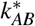 is low, both *N*_1_ and *N*_2_ are killed by *B*_1_ and *B*_2_, respectively, but as *ω*_*max*1_ *> ω*_*max*2_, *N*_1_ is killed at a faster rate, and the consortium is dominated by *N*_2_. As 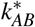 increases, both the actions of *B*_1_ and *B*_2_ are reduced so that the rate of bacteriocin caused cell death is reduced. Both *N*_1_ and *N*_2_ increase, causing the initial rise in Resveratrol production seen in the steady state manifold of Fig. 8A. As *μ*_*max*1_ *> μ*_*max*2_ and *K*_*Bmax*1_ *> K*_*Bmax*1_, the rate of population increase of *N*_1_ is faster than *N*_2_ and the quorum sensing molecule prevents the bacteriocin caused cell death of *N*_1_ more than *N*_2_. Therefore, *N*_1_ increases rapidly. Eventually, when 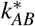 is high, both populations increase to the maximum capacity of the metabolite store, *s*, where *N*_1_ outcompetes *N*_2_, leading to a reduction and eventual extinction of *N*_2_ and no *R*, as seen on the right-hand side of the steady-state manifold of Fig. 8A. Therefore, the peak in the steady state manifold of Fig. 8A characterises the optimum rate of quorum sensing production that reduces the action of *B*_1_ and *B*_2_ enough, such that both cells can exist in large concentrations but does not allow *N*_1_ to outcompete *N*_2_ for the metabolite store, *s*. Therefore the ES can maximise the Resveratrol concentration, 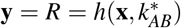, with 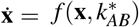 as described in Eq. (5), by optimising the cell strain population ratio. Maximising the production of Resveratrol is useful as this antioxidant is used in multiple treatments and products [50].

**Fig 7.**
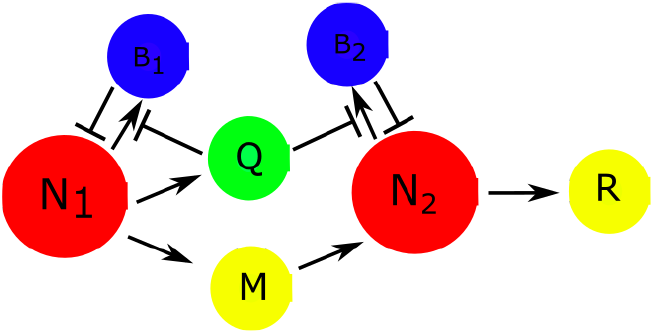
Labour Division Model - Both populations, *N*_1_ and *N*_2_ kill themselves through a bacteriocin *B*_1_ and *B*_2_, respectively, yet one population, *N*_1_, produces a quorum-sensing molecule, *Q*, which inhibits the production of both bacteriocins. A split production of Resveratrol, *R*, is included between the two strains. Parameter *k*_*AB*_ is used within the production *Q* to tune the cell population ratio and affect the production of Resveratrol, *R*.

**Fig 8.**
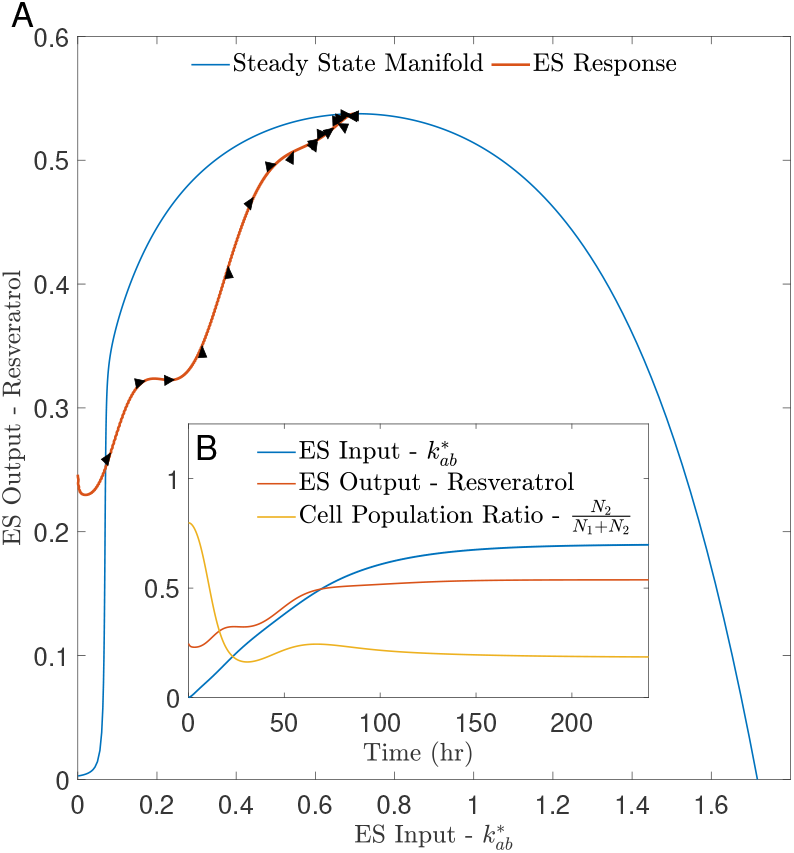
The ES controller response to the labour division ‘system to be optimised’. A) The ES response compared to the equilibrium manifold, with an arrow every ten hours in the direction of the ES response. B) The input signal, 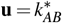, acting on the labour division system and the resulting output signal, 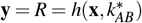. ES parameters: *k*_1_ = 0.035, *A* = 0.1 and *K* = 25.

#### 2) Simulation Results

The system in eq. (5) is used as the ‘system to be optimised’, 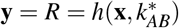, within the ES Eq. (3) to simulate the ES optimising the population ratio between the two strains to maximise the production of Resveratrol, *R*, as shown in Fig. 8. It can be seen that the embedded ES response works towards the optimum strain ratio, working towards the maximum of the Resveratrol steady-state manifold, resting on the maximum possible production of Resveratrol. Around the optimum, the ES has a similar response to Fig. 5, oscillating about the optimum, as seen by the directions of the arrows around the peak.

However, the approach to the optimum is significantly different between the static map in Fig. 5 and the labour division system in Fig. 8. The output of the ES does not follow the steady state manifold, as the time scale separation between the Labour division system and the integral/oscillator combination could not be achieved. Unfortunately, the labour division system takes roughly 10 hours for the steady state to react to a change in *k*_*AB*_. Therefore, there is no timescale separation between the ‘system to be optimised’ and the oscillator, as its time period is roughly eight minutes. This means that once the integrator has averaged out the eight minute oscillation, the ES does not react to the steady state but to some transient state of the labour division system. This can be seen in Fig. 8 as the output moves towards the optimum, it does not move along the steady-state manifold. ES still works without timescale separation, but the stability of the output cannot be guaranteed. Fortunately, biological systems are intrinsically more robust due to many factors, such as resource constraints [51].

#### 3) Sensitivity Analysis

As described in Section III-A.2, the parameters *k*_1_, *A* and *K* are chosen to tune the ES, but the maximum has to be reached via a range of parameters. Fig. 9 displays the ES system’s final steady state (not the system’s open loop to be optimised but the closed loop ES system) for a range of these parameters. For the parameter ranges 0.003 → *k*_1_ → 0.08, 0 *< A* ≤ 200 and 0 *< K* ≤ 10^*′*^000, the ES controller manages to reach the optimum.

**Fig 9.**
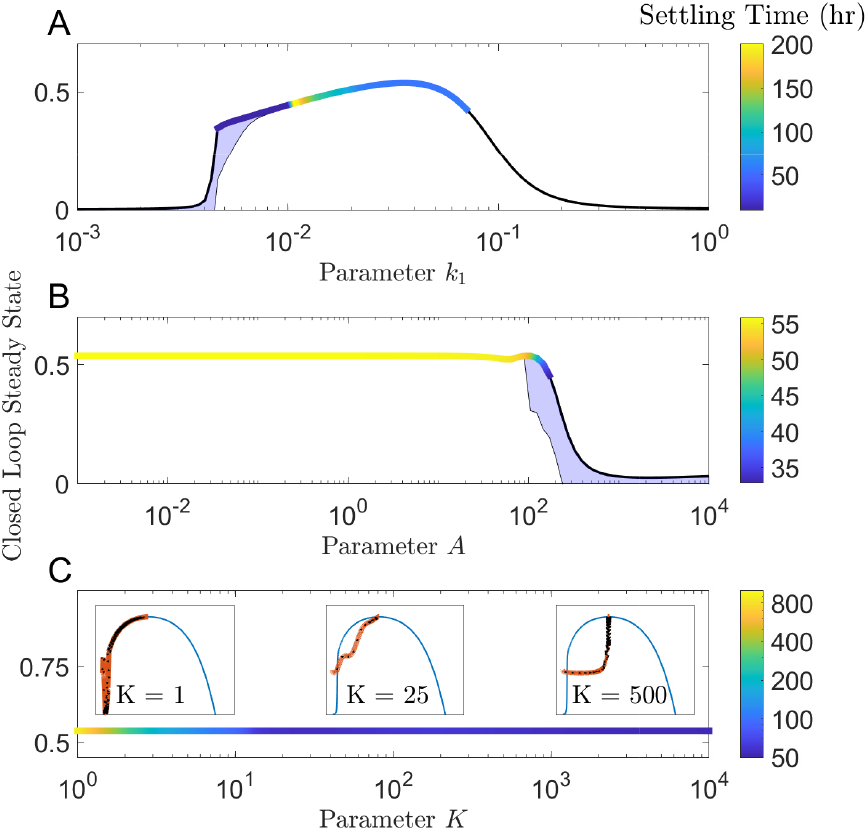
The Range of the ES parameters that achieve steady state. For each parameter, the mean steady state is surrounded by the amplitude of its steady state oscillation, which is displayed as the shaded blue area. For the range of parameters that successfully reach the optimum steady state, the colour of the mean steady-state line varies with the time taken to reach the steady state. A) The range of parameter *k*_1_. B) The range of parameter *A*. C) The range of parameter *K*, including three small panels displaying how the ES follows the equilibrium manifolds, as in Fig. 8, at points within the *K* range. ES parameters: *k*_1_ = 0.035, *A* = 0.1 and *K* = 25, unless otherwise stated.

It can be seen in Fig. 9A that outside 0.003 → *k*_1_ → 0.08, the ES fails to reach the optimum. However, it still increases the production of the Resveratrol to the range of 0.001 →*k*_1_ → 0.25. Outside 0.003 → *k*_1_ → 0.08, the ES controller requires *z*_2_ (the integrator state) to move outside its range of feasible operating concentrations, as discussed in Section. IV.

*A* controls the amplitude of the perturbation added to the input state, as can be seen in Fig. 9B. As expected, the maximum is found for all *A* values up until *A* = 200, the point at which there is no longer timescale separation between the oscillator and the integrator, as described in Section II-A. Due to the settling time of the labour division system, we do not see the same decrease in settling time, as shown in Fig. 6, due to the inadequate timescale separation between the labour division system and the oscillator/integrator combination. Although the timescale separation is not achieved, and therefore, there is no SPA stability guarantee, the ES still manages to hit the maximum.

Fig. 9C shows that *K* controls the learning rate of the controller, affecting the time to reach the maximum. *K* affects the timescale separation of the oscillator and integrator, as described in Section II-A, but when the timescale separation between the ‘system to be optimised’ and the oscillator/integrator combination cannot be achieved, the size of *K*, and the timescale separation of the oscillator and integrator, will also affect whether the system can travel along the steady state manifold of the open loop system.

As *K* is reduced, the integrator reacts slower to the observed changes in the ‘system to be optimised’, allowing more time for this system to reach steady state. Once at steady state, the slow learning rate enables the ES controller to continue to move along the steady-state manifold towards the maximum, coming closer to successful timescale separation, as can be seen in the small panel of Fig. 9C title *K* = 1. Unfortunately, for this labour division system, *K* has to be reduced to around 1 for the system to traverse the steady-state manifold from the first time the system crosses it, resulting in a settling time of 1000 hours. It can be seen from the progression of small panels in Fig. 9C that as *K* increases, the response moves further from the steady state manifold, but the settling time reduces to a lower limit of 50 hours, the settling time of the labour division when not connected to the ES. Therefore, the choice of *K* is a balance between realistic settling times and proximity to a steady state manifold as a proxy for successful timescale separation. As *K* increases, we would expect, as in Fig. 6C, that the closed loop system would become unstable and unpredictable, yet due to the natural robustness of the biological system [51], this is avoided up to higher *K* values.

#### 4) Fano Factor

It can be seen in Table. II that the Fano Factor, Eq, (4), of each ES parameter is small, showing that the controller is robust to optimisation in the presence of ES parameter uncertainty, as shown in Fig. 9. To compare the sensitivity of the ES to changes in its internal parameters to the sensitivity of Resveratrol, 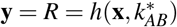, to changes in the input, the Fano Factor of the input, 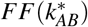, is also included in Table. II. *FF*(*k*_*AB*_) = 0.0076, three times larger than the nearest *FF* of any of the ES parameters, *FF*(*k*_1_) = 0.0025. This emphasises how insensitive the optimum is to changes in *k*_1_, *A* and *K* and the ES’s ability to adapt to changes in internal parameters and still find the optimum. However, it can be seen that the system is most susceptible to changes in *k*_1_ by three orders of magnitude. This is discussed in Section .IV.

**TABLE II.** The Fano Factor, *FF* IN EQ. (4), FOR EACH PARAMETER, *P*.

| $P$ | $k_1$ | $A$ | $K$ | $k_{AB}^*$ |
| --- | --- | --- | --- | --- |
| $FF(P)$ | 0.0025 | $1.34 \times 10^{-10}$ | $5.86 \times 10^{-6}$ | 0.0076 |

### C. Biological Implementation

The labour division system can be implemented as described in [49] with the two different Colicins [52] for the bacteriocin, *B*_1_ and *B*_2_, and acyl-homoserine lactone as a quorum sensing molecule *Q* [53], requiring the mass action kinetics described in Section III-B.1. Combined with the split production outlined in [20], adding the FjTAL gene to *N*_1_ to produce *M* and the At4CL1 and VvVST genes to *N*_2_ to turn *M* into *R*.

The ES system described in Section II-E needs to be grown in the consortium. These plasmids could be added to *N*_1_ so that the RNA that makes up *z*_2_ can be engineered to contain the sRNA component that annihilates *z*_1_ and the mRNA to promote the production of the quorum sensing molecule, *Q*. If these extra plasmids within *N*_1_ cause significant cellular stress, then the chromosomes of *N*_1_ could be edited to include the ES controller. If it were found that the dynamics of *N*_1_ were still affected by this inclusion, a third cell strain *N*_3_ could be included in the consortium, exclusively to house the ES system described in Section II-E. *N*_3_ would produce the quorum sensing molecule, *Q*, varying the population ratio of *N*_1_ and *N*_2_.

The output of the ‘system to be optimised’, *R*, needs to be sensed by ES, coactivating or corepressing *z*_1_ alongside the proteins of the oscillator, *D*_1_ and *D*_2_, respectively. To estimate the concentration of Resveratrol, the rate of expression of the final gene in the split production chain, the VvVST gene, can be coupled with the production of a quorum sensing molecule that will defuse out of *N*_2_ and can be sensed by *N*_1_ (or *N*_3_ if used). This has been proposed *in silico* [54], and used *in vitro* [55]. Different quorum sensing molecules can be used orthogonally within one system [56].

## IV. Discussion

The ES system is most sensitive to changes in *k*_1_, with a Fano Factor between three to eight magnitudes greater than *K* and *A* in both examples. *k*_1_ scales the coupling from the integrator state to the input of the system to be optimised. For the static map problem in Section III-A, the optimum input is five, using *k*_1_ around 0.25, whereas for the labour division system in Section III-B, when the optimum input is between 0.5 and 0.6 and therefore *k*_1_ must be around an order of magnitude lower at 0.035. A computational integrator can calculate the integral of an input over a large range of magnitudes as they follow a mathematical formula. Whereas the integrator used here is a living bio-synthetic system, being produced in response to the input. *k*_1_ is trying to make up for the inflexibility in the working range of the bio-synthetic integrator. Due to production rate constraints, there is a feasible range in which the bio-synthetic integrator can operate. Therefore, when using the ES, *k*_1_ will need to be tuned as the user finds this suitable range of the integrator so that it can effectively act on the ‘system to be optimised’. As the values of *K* and *A* represent pre-existing ES parameters, they do not need fine tuning, like *k*_1_, as they will only affect the controller’s performance, not its final steady state.

The parameter *ω* is part of the chosen oscillator system. This oscillator frequency, *ω*, must be selected to balance the timescale separation between the ‘system to be optimised’ and the integrator.

The parameter *A* scales the perturbations in the input; this can be seen in Fig. 5A and Fig. 8A. The size of *A* affects the parameter space in which the ES explores each perturbation. The larger the *A*, the more likely the system is to become unstable due to a mismatch between the integrals estimate of the average system and the actual system.

Finally, *K* implements a convolution of the effect of the input on the system to be optimised alongside a scaled perturbation that was added to the input, allowing the integrator to estimate the average of the system. This is the learning rate of the ES. The larger *K*, the faster the ES will reach a steady state; however, a larger *K* could lead to instability. When a biological system is added to the ES, *K* can be used to control the success of timescale separation, tuning how close the system moves along the steady state manifold. It can be seen that for the labour division system, in Fig. 9C, even with a large *K*, the system still manages to reach the optimum, as the tuning of *k*_*AB*_, has the same effect on Resveratrol production during the systems transient response, and during its steady state. Therefore, *K* must be chosen carefully to achieve a maximum in an adequate time without causing instability.

In summary, the parameters *ω, K* and *A* are equivalent to the frequency, learning rate and perturbation gain, respectively, usually found in an extremum-seeking controller’s [19] with the stability characteristics described in [30]. *k*_1_ is an added ES parameter needed to scale the bio-synthetic integral system as it acts on the ‘system to be optimised’.

## V. Conclusion

Through this work, it has been shown that an extremum-seeking controller could be implemented with a molecular network using synthetic biology. This ES could be coded into bacteria, allowing them to optimise cellular processes within their own culture.

Extremum seeking does not have stability guarantees if time scale separation is not ensured. This is tricky for biological systems as restricted systems exist that can be used for the oscillator and integrator. Even with this limitation, the ES system can still reach a steady state, optimising some internal elements of the system to maximise the output.

This work is implemented *in-silico* and should be tested *in vitro* to grow the first example of a biological optimiser. Each element of the ES is modular and, therefore, can be implemented in parts. Initially, *in vitro*, the oscillations could be implemented via external perturbations on the bioreactor, [57], to simplify the system. Then, a biological oscillator can be implemented in its place (such as the Repressilator, which is used as an example here). This embedded ES can then be implemented into many different biological systems, such as the labour division system, to form a biological optimiser.

## Author Contributions

B.S. designed and carried out the simulations in Matlab2024a and wrote the manuscript with inputs from all the authors. José I. Jiménez assisted in our search for biological maximisation problems and the biological implementation of the systems. L.M. and L.R. conceived the project, contributed to manuscript writing, and supervised the entire work.

## Acknowledgement

An EPSRC DTP Scholarship (grant EP/T51780X/1) supports B.S. L.M. is supported by an EPSRC fellowship (grant EP/S01876X/1) and a BBSRC transition award (grant BB/W013959/1).

## Data Availability

Data sharing does not apply to this article as no datasets were generated or analysed during the current study.

## Code Availability

The code for this study can be accessed via this link https://github.com/Ben-Smart/Extremum_Seeking_LD.

## References

[1] M. Khammash, M. Di Bernardo, and D. Di Bernardo. Cybergenetics: Theory and Methods for Genetic Control System. Proceedings of the IEEE Conference on Decision and Control, 2019-December:916–926, 12 2019.

[2] Mustafa H. Khammash. Cybergenetics: Theory and Applications of Genetic Control Systems. Proceedings of the IEEE, 110(5):631–658, 5 2022.

[3] Marco P.C. Marques and Pedro Fernandes. Microfluidic Devices: Useful Tools for Bioprocess Intensification. Molecules 2011, Vol. 16, Pages 8368-8401, 16(10):8368–8401, 9 2011.

[4] Sant Kumar and Mustafa Khammash. Platforms for Optogenetic Stimulation and Feedback Control. Frontiers in Bioengineering and Biotechnology, 10:918917, 6 2022.

[5] Aleksey Malyshev et al. Advantages and Limitations of the Use of Optogenetic Approach in Studying Fast-Scale Spike Encoding. PLoS ONE, 10(4):e0122286, 4 2015.

[6] Wolfgang Halter, Simon Michalowsky, and Frank Allgower. Extremum seeking for optimal enzyme production under cellular fitness constraints. 2019 18th European Control Conference, ECC 2019, pages 2159–2164, 6 2019.

[7] M. Atkinson et al. Development of Genetic Circuitry Exhibiting Toggle Switch or Oscillatory Behavior in Escherichia coli. Cell, 113(5):597–607, 5 2003.

[8] J. Stricker et al. A fast, robust and tunable synthetic gene oscillator. Nature 2008 456:7221, 456(7221):516–519, 10 2008.

[9] T. Sohka et al. An externally tunable bacterial band-pass filter. Proceedings of the National Academy of Sciences of the United States of America, 106(25):10135–10140, 6 2009.

[10] Uri Alon. An Introduction to Systems Biology. An Introduction to Systems Biology, 7 2019.

[11] N. Rosenfeld et al. Negative Autoregulation Speeds the Response Times of Transcription Networks. Journal of Molecular Biology, 323(5):785–793, 11 2002.

[12] A. Harris et al. Designing Genetic Feedback Controllers. IEEE Transactions on Biomedical Circuits and Systems, 9(4):475–484, 8 2015.

[13] S. Aoki et al. A universal biomolecular integral feedback controller for robust perfect adaptation. Nature 2019 570:7762, 570(7762):533–537, 6 2019.

[14] Y. Qian et al. Realizing ‘integral control’ in living cells: how to overcome leaky integration due to dilution? Journal of The Royal Society Interface, 15(139), 2018.

[15] N. Olsman et al. Architectural Principles for Characterizing the Performance of Antithetic Integral Feedback Networks. iScience, 14:277–291, 4 2019.

[16] Michael Chevalier et al. Design and Analysis of a Proportional-Integral-Derivative Controller with Biological Molecules. Cell Systems, 9(4):338–353, 10 2019.

[17] Max Whitby et al. PID Control of Biochemical Reaction Networks. IEEE Transactions on Automatic Control, 67(2):1023–1030, 2 2022.

[18] Michael B. Elowitz and Stanislas Leibier. A synthetic oscillatory network of transcriptional regulators. Nature 2000 403:6767, 403(6767):335–338, 1 2000.

[19] Y. Tan et al. Extremum Seeking From 1922 To 2010. Technical report, Proceedings of the 29th Chinese Control Conference, Beijing, 7 2010.

[20] Huadong Peng et al. A molecular toolkit of cross-feeding strains for engineering synthetic yeast communities. Nature Microbiology 2024 9:3, 9(3):848–863, 2 2024.

[21] C. Mayhew et al. Robust source-seeking hybrid controllers for autonomous vehicles. Proceedings of the American Control Conference, pages 1185–1190, 2007.

[22] J. Beaudoin et al. Bluff-body drag reduction by extremum-seeking control. Journal of Fluids and Structures, 22(6-7):973–978, 8 2006.

[23] I. Haskara et al. Multivariable EGR/spark timing control for IC engines via extremum seeking. Proceedings of the American Control Conference, 2006:1173–1178, 2006.

[24] M. Krstic et al. Multivariable adaptive control of instabilities arising in jet engines. Control Engineering Practice, 14(7):833–842, 7 2006.

[25] E. Schuster et al. Optimal beam matching in particle accelerators via extremum seeking. Proceedings of the American Control Conference, 2006:1962–1967, 2006.

[26] H. Wang et al. Optimizing bioreactors by extremum seeking. International Journal of Adaptive Control and Signal Processing, 13(8):651–669, 12 1999.

[27] G. Bastin et al. On extremum seeking in bioprocesses with multivalued cost functions. Biotechnology Progress, 25(3):683–689, 5 2009.

[28] M. Guay et al. Adaptive extremum seeking control of continuous stirred tank bioreactors with unknown growth kinetics. Automatica, 40(5):881–888, 5 2004.

[29] Y. Tan et al. On the choice of dither in extremum seeking systems: A case study. Automatica, 44(5):1446–1450, 5 2008.

[30] Ying Tan, Dragan Nešić, and Iven Mareels. On stability properties of a simple extremum seeking scheme. Proceedings of the IEEE Conference on Decision and Control, pages 2807–2812, 2006.

[31] James Murdock, Jan Sanders, and Ferdinand Verhulst. Averaging methods in nonlinear dynamical systems. Applied Mathematical Sciences (Switzerland), 59:i–431, 2007.

[32] Alexander Scheinker, En Chuan Huang, and Charles Taylor. Extremum Seeking-Based Control System for Particle Accelerator Beam Loss Minimization. IEEE Transactions on Control Systems Technology, 30(5):2261–2268, 9 2022.

[33] Alexander Scheinker. Application of Extremum Seeking for Time-Varying Systems to Resonance Control of RF Cavities. IEEE Transactions on Control Systems Technology, 25(4):1521–1528, 7 2017.

[34] Mouhacine Benosman and Gökhan M. Atnç. Extremum seeking-based adaptive control for electromagnetic actuators. International Journal of Control, 88(3):517–530, 3 2015.

[35] Jan Maximilian Montenbruck et al. Extremum Seeking with Drift IFAC-PapersOnLine, 48(11):126–130, 1 2015.

[36] Alexander Scheinker. Simultaneous stabilization and optimization of unknown, time-varying systems. Proceedings of the American Control Conference, pages 2637–2642, 2013.

[37] C. Briat et al. Antithetic Integral Feedback Ensures Robust Perfect Adaptation in Noisy Bimolecular Networks. Cell Systems, 2(1):15–26, 1 2016.

[38] Béla Novák and John J. Tyson. Design principles of biochemical oscillators. Nature Reviews Molecular Cell Biology 2008 9:12, 9(12):981–991, 10 2008.

[39] J. Hasty et al. Synthetic Gene Network for Entraining and Amplifying Cellular Oscillations. Physical Review Letters, 88(14):148101, 3 2002.

[40] Dan Siegal-Gaskins et al. Gene circuit performance characterization and resource usage in a cell-free “breadboard”. ACS Synthetic Biology, 3(6):416–425, 6 2014.

[41] Wolfgang Halter et al. A resource dependent protein synthesis model for evaluating synthetic circuits. Journal of Theoretical Biology, 420:267–278, 5 2017.

[42] Wolfgang Halter, Richard M. Murray, and Frank Allgöwer. Analysis of primitive genetic interactions for the design of a genetic signal differentiator. Synthetic Biology, 4(1), 1 2019.

[43] B. Ingalls. Mathematical modeling in systems biology : an introduction. The MIT Press, 2013.

[44] W. G. Spiegelman et al. Bidirectional Transcription and the Regulation of Phage λ Repressor Synthesis. Proceedings of the National Academy of Sciences, 69(11):3156–3160, 11 1972.

[45] Hsin Ho Huang, Yili Qian, and Domitilla Del Vecchio. A quasi-integral controller for adaptation of genetic modules to variable ribosome demand. Nature Communications 2018 9:1, 9(1):1–12, 12 2018.

[46] David K. Summers. The kinetics of plasmid loss. Trends in Biotechnology, 9(1):273–278, 1 1991.

[47] Andras Gyorgy et al. Isocost Lines Describe the Cellular Economy of Genetic Circuits. Biophysical Journal, 109(3):639–646, 8 2015.

[48] Tim A. Hoek et al. Resource Availability Modulates the Cooperative and Competitive Nature of a Microbial Cross-Feeding Mutualism. PLOS Biology, 14(8):e1002540, 8 2016.

[49] Behzad D. Karkaria, Alex J.H. Fedorec, and Chris P. Barnes. Automated design of synthetic microbial communities. Nature Communications 2021 12:1, 12(1):1–12, 1 2021.

[50] Mohammadreza Rostami et al. Resveratrol-loaded nanocarriers: Characteristics, sources, health effects, recent delivery systems, and their food and biomedical applications. Food Bioscience, 61:104845, 10 2024.

[51] Hiroaki Kitano. Towards a theory of biological robustness. Molecular Systems Biology, 3, 9 2007.

[52] Volkmar Braun, Silke I. Patzer, and Klaus Hantke. Ton-dependent colicins and microcins: modular design and evolution. Biochimie, 84(5-6):365–380, 5 2002.

[53] Mair E.A. Churchill, Hiruy M. Sibhatu, and Charis L. Uhlson. Defining the Structure and Function of Acyl-Homoserine Lactone Autoinducers. Methods in Molecular Biology, 692:159–171, 2011.

[54] Vittoria Martinelli et al. Multicellular PI Control for Gene Regulation in Microbial Consortia. IEEE Control Systems Letters, 6:3373–3378, 2022.

[55] Ye Chen et al. Emergent genetic oscillations in a synthetic microbial consortium. Science, 349(6251):986–989, 8 2015.

[56] Stefan J. Tekel et al. Engineered orthogonal quorum sensing systems for synthetic gene regulation in escherichia coli. Frontiers in Bioengineering and Biotechnology, 7(MAR):444033, 4 2019.

[57] Peter L.. Silveston and R. R. Hudgins. Periodic operation of reactors. page 793, 2013.

